# The effect of dopamine D2 receptor blockade on human motor skill learning

**DOI:** 10.1101/2023.09.01.556003

**Authors:** Eleanor M. Taylor, Dylan Curtin, Trevor T-J. Chong, Mark A. Bellgrove, James P. Coxon

## Abstract

*Rationale:* Dopamine signalling supports motor skill learning in a variety of ways, including through an effect on cortical and striatal plasticity. One neuromodulator that has been consistently linked to motor skill learning is dopamine. However, the specific role of dopamine D2 receptor in the acquisition and consolidation stages of motor learning remains unclear.

**Objectives:** To examine the effect of a selective D2 receptor antagonist on human motor skill acquisition and consolidation.

**Methods:** In this randomised, double-blind, placebo-controlled design, healthy adult men and women (N = 23) completed a sequential motor skill learning task after taking either sulpiride (800mg) or placebo. A 20-minute bout of high-intensity interval cycling exercise was included to enhance experimental effects and counteract potentially confounding sedative effects of sulpiride.

**Results:** Sulpiride reduced performance during motor skill acquisition relative to placebo in the first session, however this difference was abolished at the subsequent retention test. Sulpiride did not reduce consolidation of learning as expected, however it led to a reduction in speed of execution relative to placebo.

**Conclusions:** Our results demonstrate that neuromodulation at the dopamine D2 receptor is critical in the early acquisition of a novel motor skill. These results may have functional relevance in motor rehabilitation as reduced dopamine transmission can impact performance during initial learning and slow subsequent performance of the skill.

## Introduction

Motor skill learning is the process by which task performance becomes more automatic and efficient with practice (Krakauer et al. 2019), and is a key aspect of motor rehabilitation (Krakauer 2006). It includes online acquisition, when the task is actively practiced and improved, and offline consolidation, whereby the skill is encoded into memory and becomes resistant to interference (Dayan and Cohen 2011). Dopaminergic signalling in the basal ganglia plays a key role in motor learning, facilitating coordinated motor output and modulating reward and error signals associated with a given movement (Coddington and Dudman 2019; Robbins and Everitt 1992; Schultz 2015).

Dopamine projections from the substantia nigra primarily exert effects on the excitatory ‘direct’ and inhibitory ‘indirect’ basal ganglia pathways, modulated by the dopamine D1 and D2 receptors, respectively (Albin et al. 1989; DeLong et al. 1986; Dreyer et al. 2010). It is thought that coordinated signalling along both direct and indirect pathways is required to support motor learning (Wiecki and Frank 2010). Dopamine also supports motor learning by modulating cortical plasticity.

Rodent studies have shown that disrupting dopamine signalling to the primary motor cortex (M1) limits the induction of plasticity (Hosp et al. 2009; Molina-Luna et al. 2009), which is fundamental in consolidating a newly learned motor skill (Hosp et al. 2011; Rioult-Pedotti et al. 2015). Studies in mice suggest early motor learning primarily relies on D1 receptor activity (Nakamura et al. 2014; Sommer et al. 2014), with increasing reliance on dopamine D2 signalling in the later stages of motor learning, as the task becomes consolidated and more automatic (Henry et al. 2009; Sommer et al. 2014).

There is also indirect evidence supporting the importance of dopamine D2 activity in motor skill learning in humans. Patients with Parkinson’s disease (PD), an age-related disorder characterised by progressive loss of dopamine neurons, demonstrate reduced capacity to modulate plasticity in M1 (Kishore et al. 2012; Suppa et al. 2011; Ueki et al. 2006), which is associated with deficits in motor learning (Marinelli et al. 2017; Ueda et al. 2022). In healthy populations, online motor learning is modulated by genetic variations associated with dopamine receptor expression (Noohi et al. 2016; Schuck et al. 2013). Specifically, Noohi and colleagues (2014) found that genetic variations linked to D2 receptor availability were associated with motor sequence learning, but not visuomotor adaptation. Similarly, Beatu and colleagues (2015) found that gene variations characterised by greater D2 receptor availability were associated with improved recovery following interference. Together, these studies provide correlational evidence for the importance of dopamine transmission involving the D2 receptor in human motor learning.

Pharmacological challenge studies allow for more direct examination of the effects of dopamine in motor learning in humans. Blockade of dopamine via the selective D2 antagonist, sulpiride, reduces long-term potentiation (LTP)-like plasticity in M1 in healthy adults (Monte-Silva et al. 2011). Further, a study by Meintzschel and Ziemann (2006) demonstrated that use-dependent plasticity within the motor cortex of healthy adults was reduced by the non-selective D2 antagonist haloperidol. Another study by Floel and colleagues (2005) investigated the effect of increasing dopamine levels with levodopa on a simple thumb abduction task. Results showed levodopa increased thumb accelerations in younger adults, and improved thumb acceleration learning in older adults. Importantly, this task required minimal cognitive effort. Pharmacological increases in dopamine may improve some aspects of motor learning, particularly in cases of dopamine degeneration, however it can also negatively impact cognitive function, including motor sequence learning (Vaillancourt et al. 2013). Taken together, these findings show that dopamine activity supports processes associated with motor learning, however the causal role of the D2 receptor in different stages of motor skill learning requires further direct examination in humans.

Acute exercise, purported to support endogenous upregulation of dopamine transmission (Hattori et al. 1994; Sacheli et al. 2019), has also been shown to improve motor learning (Wanner et al. 2020). Furthermore, improvements in offline learning after an acute bout of exercise have been associated with genetic variations in D2 dopamine availability (Christiansen et al. 2019; Mang et al. 2017). We have recently shown that the selective D2 antagonist, sulpiride, attenuates the modulation of cortical excitation-inhibition balance induced by exercise (Curtin et al. 2023). This reseach implicates the D2 receptor in exercise-enhanced plasticity and learning, however causal evidence is needed.

In the present study, we aimed to investigate the role of the D2 receptor on human motor skill learning following exercise. We used the D2 receptor antagonist sulpiride, which acts predominantly on post-synaptic receptors at an 800 mg dose, with a low affinity for D1 and non-dopamine receptors (Miyamoto et al. 2005). We included a bout of high intensity exercise prior to motor learning, and utilised a continuous motor sequence task, allowing task complexity to reflect real-world learning (Luft and Buitrago 2005; Reis et al. 2009). Exercise has been shown to improve motor learning on this task (Stavrinos and Coxon 2017) and was expected to enhance experimental effects by supporting motor learning in the placebo condition. Additionally, Exercise was intended to increase alertness and arousal (Dietrich and Audiffren 2011) and counteract the known mild sedative effects of sulpiride (Ho et al. 2009; Loy et al. 2013) which may obscure any apparent effects on learning. Given the specificity of sulpiride to D2-like receptors, and evidence that early learning relies primarily on D1 receptor activity, participants in both sulpiride and placebo conditions were expected to demonstrate motor skill learning during online acquisition. We hypothesised that sulpiride would reduce consolidation of a new motor skill, with reduced offline consolidation following exercise for sulpiride compared to placebo.

## Methods

### Participants

Twenty-three healthy young adults participated (11 female, mean age = 24.14 ± 3.93 years, range 19 – 34 years). Prior to the study, participants were screened for psychiatric conditions using the Mini International Neuropsychiatric Interview (Lecrubier et al. 1997) and contraindications to exercise (Adult Pre-Exercise Screening System; Norton 2012) or the study drug. Exclusion criteria included a history of psychiatric or neurological illness, significant drug or alcohol use, and the use of psychoactive drugs or medications within the last 6 months. Hormonal contraception may impact dopamine function (Algeri et al. 1976; Taylor et al. 2022), therefore female participants were excluded if not using the oral contraceptive pill or hormonal implant. One participant was left-handed, as determined by the Edinburgh Handedness Inventory (Oldfield 1971). All participants provided informed consent and were reimbursed for their time. The study was conducted in accordance with the Declaration of Helsinki and approved by the Monash University Human Research Ethics Committee.

### Design

The present study utilized a randomized, double-blind, placebo-controlled, within-participants design. Participants attended three sessions in total, comprising two testing sessions and a follow-up session (Figure 1A). During each testing session, participants ingested a capsule containing either 800 mg sulpiride or a placebo, completed a 20-minute bout of high intensity exercise, and trained in one version of the motor learning task (either SVIPT_A_ or SVIPT_B_). The order of drug and placebo was double blinded, and task version and drug order were randomised and counterbalanced. An independent researcher not involved in recruitment, data collection, or analysis oversaw the preparation and randomisation of the study drug. The exercise utilised a 20-minute high-intensity interval protocol as outlined in Curtin et al. (2023), see Supplementary Material for further detail. Participants were asked to refrain from strenuous physical activity in the 24 hours prior to each session and wear a Polar H10 heart rate monitor (Polar Electro, Finland) during testing sessions. Retention of each motor learning task was assessed approximately 7 days after initial training.

**Fig. 1.**
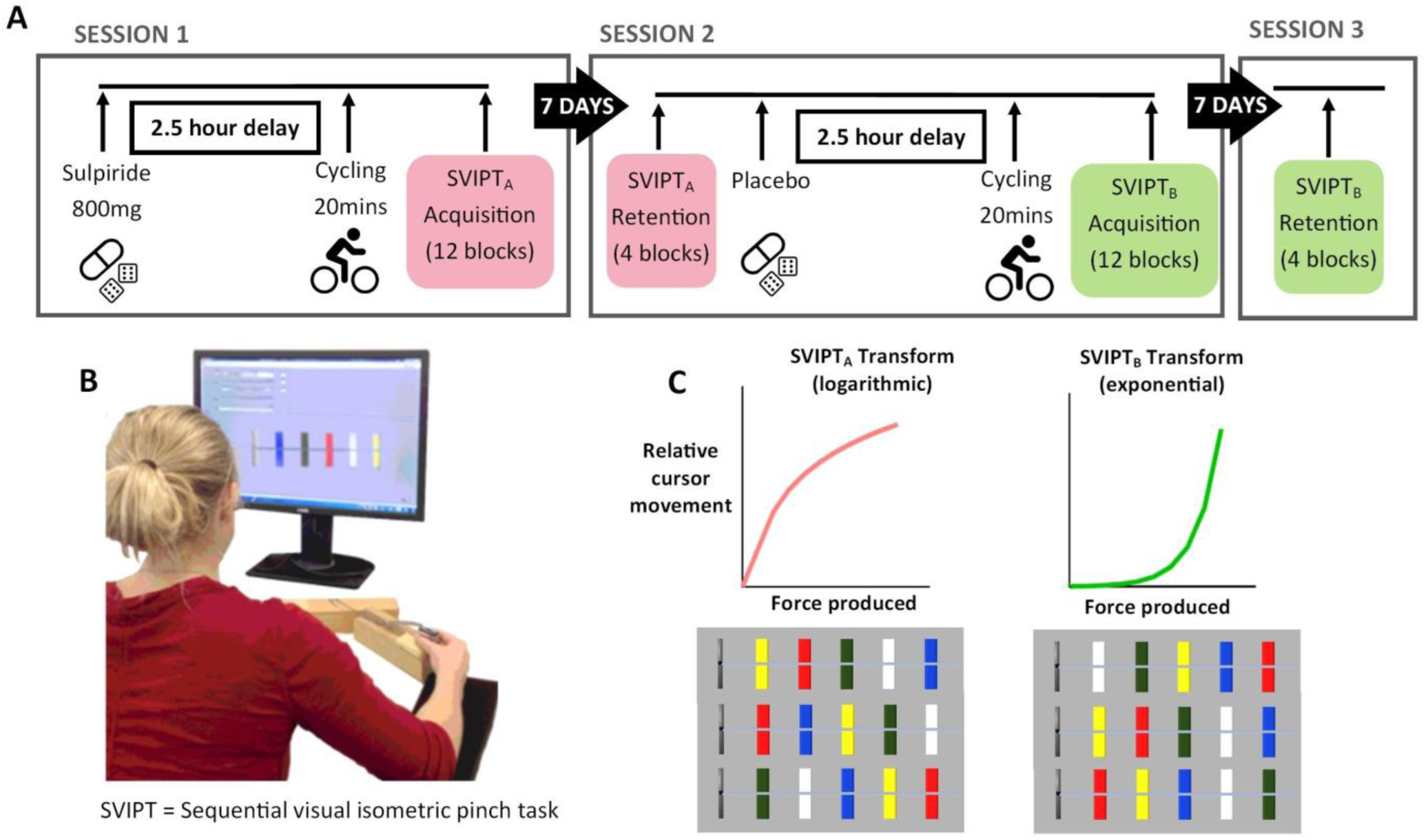
Overview of the study design. (A) Overview of testing schedule. Sessions 1 and 2 were learning sessions, involviong ingestion of the relevant capsule, 20 minutes of high-intensity interval cycling, and 12 blocks of a motor learning task. During each of these sessions, participant wellbeing and alertness was assessed hourly using Visual Analogue Scales (VAS). Retention for each motor learning task was assessed 7 days after initial learning. Note: drug and task conditions were counterbalanced across participant and learning session. (B) Depiction of SVIPT motor task adapted from Stavrinos & Coxon (2017) (C) Representation of transformations applied to the relationship between force produced and cursor movement for SVIPTA and SVIPTB and the three target orders used for individual trials necessitating the learning and trial-wise preparation of different motor execution sequences. The versions of the task had a distinct transformation applied to the relationship between force produced and relative cursor movement as well as distinct motor sequence requirements across learning sessions to ensure adequate task difficulty

### Pharmacological intervention

Participants ingested 800 mg sulpiride or an equivalent placebo capsule (microcrystalline cellulose) 2.5 hours prior to the exercise protocol to ensure peak sulpiride plasma concentration during exercise and motor learning (Caley and Weber 1995). This dose was selected to ensure adequate D2 receptor occupancy (Takano et al. 2006) and has been used previously with healthy controls without significant adverse effects (Naef et al. 2017). Participant wellbeing and alertness was assessed each hour using Visual Analogue Scales (VAS) (Bond and Lader 1974).

### Measures

#### Motor Learning Task

Participants practiced a sequential visual isometric pinch task (SVIPT), a sequential motor learning task, similar to that described by Stavrinos & Coxon (Stavrinos and Coxon 2017). Participants were seated before a computer and held a force transducer between the thumb and index finger of their dominant hand. Squeezing the force transducer produced a proportional movement of a cursor. On each trial, participants were presented with five coloured targets located along a horizontal axis (Figure 1B). They were instructed to produce five pulses of force to move the cursor to each target as quickly and accurately as possible according to a specified colour sequence (red-blue-green-yellow-white). Target locations were pseudorandomly shuffled among 3 different orders, requiring the learning and preparation of 3 different motor execution sequences in each learning session. The amount of force required to reach the furthest target was set at 45% of each participants’ maximum voluntary pinch contraction (MVC).

This study used two variations of the SVIPT task to ensure novel motor skill learning occurred across learning sessions. SVIPT_A_ applied a logarithmic transformation to the relationship between pinch force and cursor movement (see Reis et al. 2009), whereas SVIPT_B_ applied an exponential transformation to this relationship (see Cantarero et al. 2013). This varied the force output required to reach the targets. In addition, the 3 motor execution sequences that were practiced in SVIPT_A_ were different from those practiced in SVIPT_B_ (Figure 1C). Participants were not informed of this manipulation.

For both tasks, participants completed 9 familiarisation trials. They then completed 12 blocks of 12 trials, which comprised the motor skill acquisition phase. Participants were shown a visual representation of their calculated skill level after each block, to assess their progress and to encourage improvement. The delayed retention test comprised a warm-up (6 trials of the same task which were not analysed) to counteract the established warm-up decrement for this task (Reis et al. 2015) followed by four blocks of 12 trials to assess skill retention.

### Data Analysis

Performance on the SVIPT was assessed based on mean accuracy and speed for each block of trials. Trial accuracy was the summed distance from each of the five targets to their respective force peaks, resulting in a force error score, with lower force error indicating greater accuracy. The speed of each trial was calculated as the duration from trial onset to the end of the final force peak.

One participant was excluded due to failure to complete all acquisition blocks. For the remaining sample, 93/8448 trials (1.1%) were identified as outliers (*z*= >±3.29) and were winsorized to eliminate undue bias prior to calculating block means for each participant. Two participants were excluded from the retention analysis due to significantly delayed retention tests (>30 days, due to COVID-19) resulting in N = 20 for retention analyses.

To assess motor skill learning, an overall skill measure was calculated for each block that accounts for the known trade-off between speed and accuracy during motor performance, per the procedure outlined by Reis et al. (2009) and Stavrinos and Coxon (2017). The speed-accuracy trade-off function for SVIPT_A_ has previously been defined as:

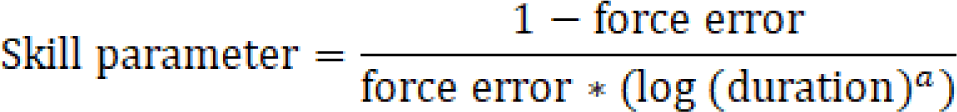

Where duration refers to the mean trial time for the block, and value of *a* is 1.627 (Stavrinos and Coxon, 2017). The same formula was applied to capture the speed-accuracy trade-off function for SVIPT_B_ (Cantarero et al., 2013). The skill measure used in analysis was the logarithm of this skill parameter, to ensure homogeneity of variance across participants (Reis et al., 2009).

### Statistical Analysis

Baseline performance was assessed using a 2 × 2 mixed ANOVA, with Session (session 1, session 2) as a within-subjects factor, and Drug Order (placebo first, sulpiride first) as a between-subjects factor.

A linear mixed model (LMM) was constructed to evaluate differences in skill between sulpiride and placebo conditions across the 12 blocks of motor skill acquisition. For a detailed description of model fitting, see Supplementary Material. The selected models included Drug Condition (placebo, sulpiride), Block Number (1 - 12) and Session (1, 2) as fixed effects, with all interaction terms included. Participant was included as a random factor, with a random slope for Block Number (Table 1). Differences were followed-up using Bonferroni-adjusted pairwise comparisons of model estimates and trends. Additional models were constructed using the above criteria to assess force error and trial time.

**Table 1.**
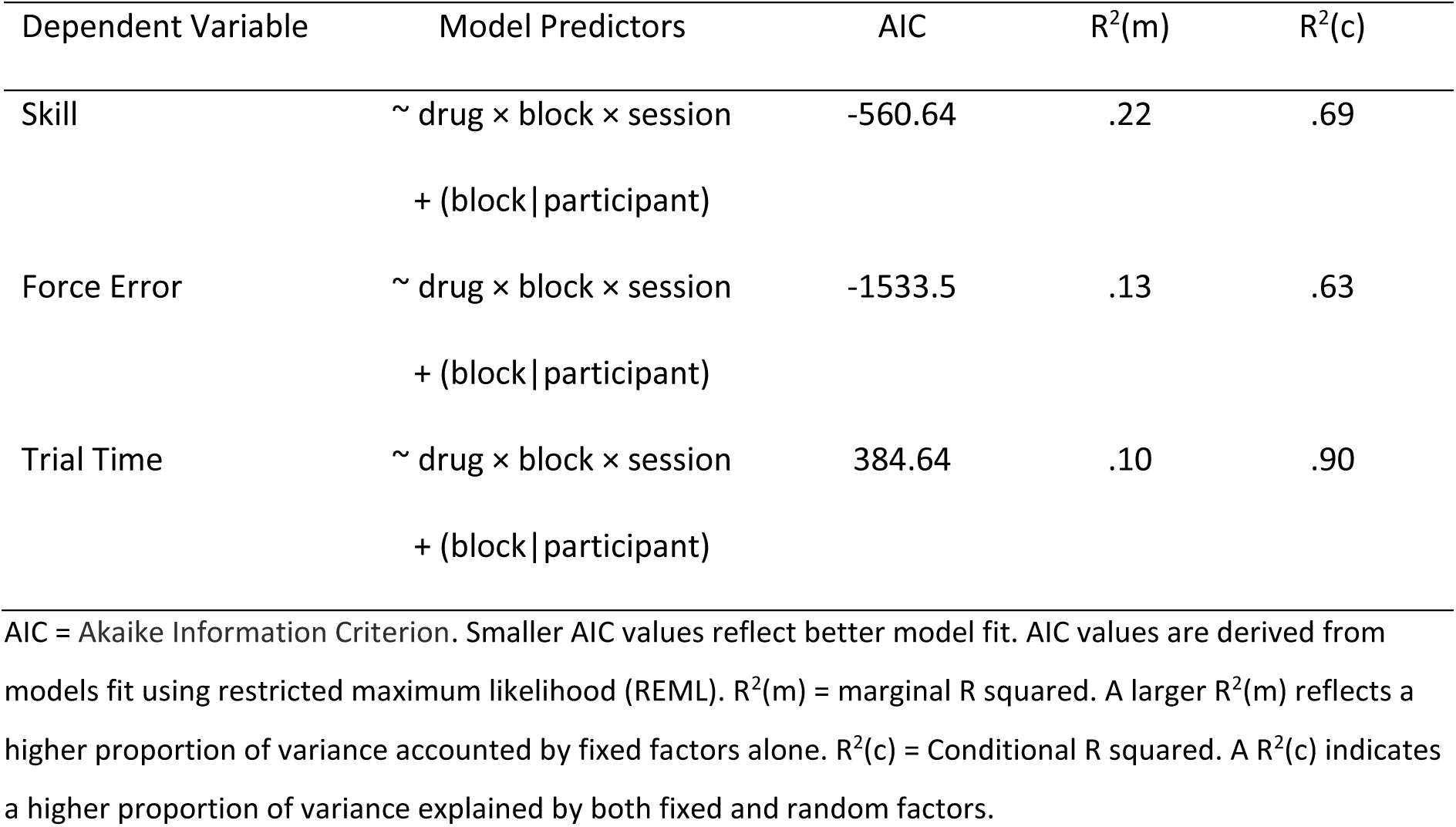
Summary of Linear Mixed Models.

Retention of motor learning was assessed by calculating the difference in performance between the end of learning (average across blocks 11 and 12) and the beginning of the retention test (average across blocks 13 and 14). Retention tests took place approximately 7 days after each learning session (Placebo, *M* = 7.75 days, *SD* = 2.22; Sulpiride, *M =* 7.10 days, *SD* = 0.44). A spearman correlation was conducted to assess whether retention test delay was correlated with performance. A 2×2 Mixed ANOVA was conducted with Learning Session (acquisition completed in session 1 or session 2) as a within-subjects factor, and Drug Order (placebo first, sulpiride first) as a between-subjects factor. Significant effects were followed-up using Bonferroni-adjusted pairwise comparisons. Effect sizes are represented by beta estimates and 95% confidence intervals. Results are reported as mean ± standard deviation, or model estimates and 95% confidence intervals. α was set to .05 for all analyses.

## Results

### Baseline

Participants’ MVC did not differ across placebo (*M=* 45.55 newtons*, SD=*13.55) and sulpiride (*M=*49.68 newtons*, SD=*14.43) conditions (*t*(1, 21) = -1.46, *p* =.16). Subjective calmness ratings were lower immediately following exercise compared to other time points, however this did not differ significantly between placebo and sulpiride sessions, and likely reflects the anxiolytic effects of high intensity exercise (see Supplementary Materials for a full summary). Examination of block 1 performance for both sessions showed skill was higher at baseline for session 2 compared to session 1 (*F*(1, 20) = 9.54, *p* = .006, 𝜂_p_^2^ = .32) reflecting familiarity with general task features. Importantly, baseline performance did not differ across Drug Order (*F*(1, 20) = 0.14, *p* = .71), nor was there a Drug Order by Session interaction (*F*(1, 20) = 0.11, *p* = .74). Mean scores for skill, force error and trial time across all blocks are illustrated in Figure 2.

**Fig. 2.**
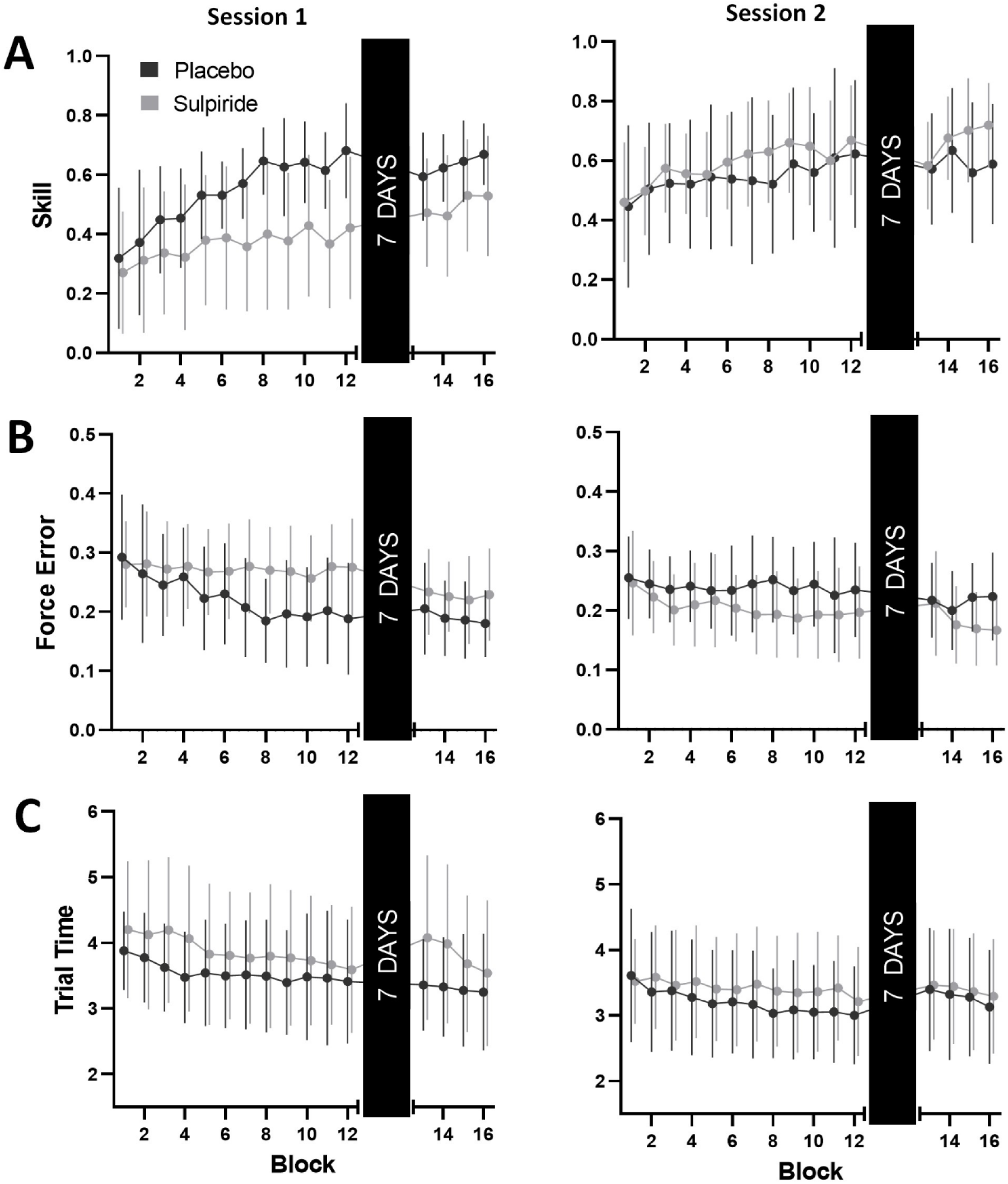
Summary of SVIPT Performance by Session and Drug Condition. Data are presented as observed means ± standard error. (A) Mean skill by Drug Condition (Black circles, Placebo; Grey circles, Sulpiride) and Learning Session (Left panel, Session 1; Right panel, Session 2). (B - C) Force error and trial time subcomponents of skill measure by Drug Condition and Session. Lower force error and trial time reflect higher skill

**Fig. 3.**
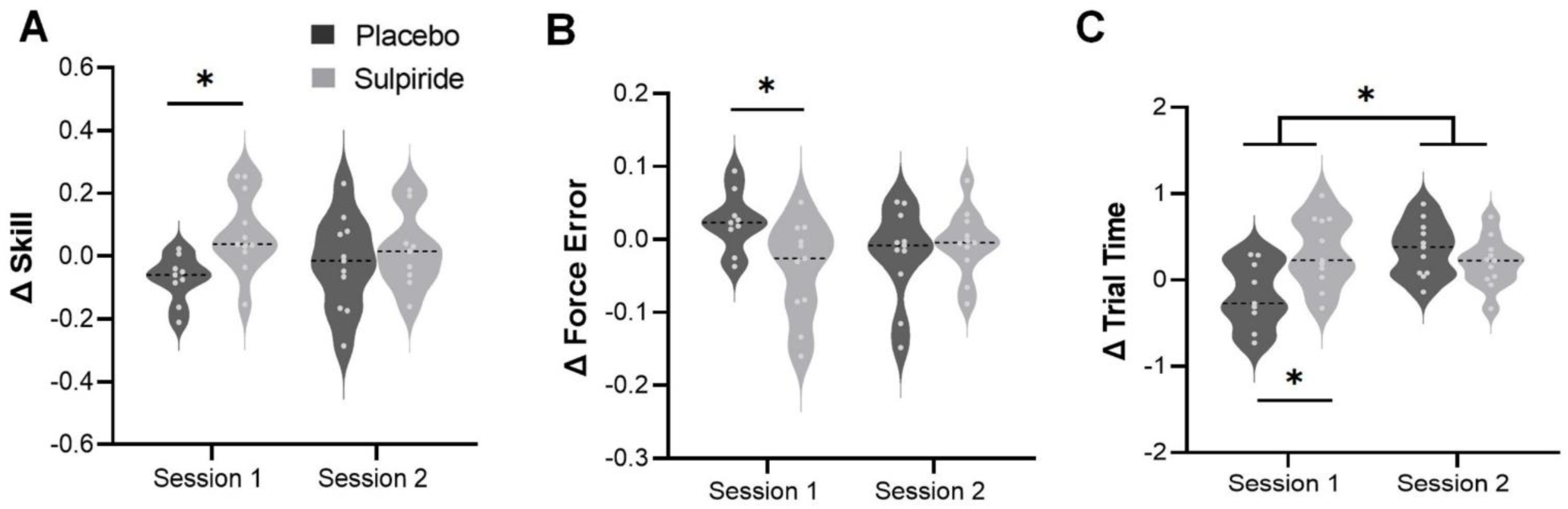
Change scores for skill, error and trial time measured at the retention test. Change scores calculated based on performance at the end of learning (session 1 or 2) and performance at the start of the subsequent retention test 1 week later. (A) Participants taking sulpiride in session 1 showed improved offline skill retention compared to placebo, with no difference between sulpiride and placebo in session 2. (B) Participants taking sulpiride in session 1 showed offline improvements in accuracy, represented by a negative error score, compared to placebo. (C) Participants taking placebo in session 1 showed an offline increase in speed at the retention test, represented by a negative trial time score, compared to sulpiride. Participants in both groups showed a decrease in speed following session 2. \**p* < .05

### Acquisition

Analysis of skill scores revealed no main effect of Drug Condition (*F*(1, 477.59) = 0.031, *p* = .58), though there was a main effect of Block (*F*(1, 20.10) = 74.84, *p* <.001), indicating an increase in skill across practice (^𝛽^^ = 0.03, 95% CI [0.02, 0.04]). A full summary of main effects, interactions and fixed effects estimates for all LMMs can be found in the Supplementary Materials, but most notably there was a three-way interaction for Drug Condition, Block and Session (*F*(1, 20.10) = 74.84, *p* =.01). During session 1, placebo participants showed greater improvements in skill across blocks compared to sulpiride (^𝛽^^ = -0.002, 95% CI [0.004, 0.02], *p* < .001). However, this difference was not evident in session 2 (^𝛽^^ = 0.01, 95% CI [-0.01, 0.004], *p* = .41). Differences in skill were driven by changes in the accuracy subcomponent of the skill measure. Force error scores showed no main effect of Drug (*F*(1, 477.32) = 1.60, *p* = .21), however there was once again a main effect of Block (*F*(1, 19.95) = 14.14, *p* = .001), and a three-way interaction for Drug Condition, Block and Session (*F*(1, 19.95) = 8.13, *p* = .01). Placebo showed greater reductions in force error across blocks compared to sulpiride in Session 1 (^𝛽^^ = -0.004, 95% CI [-0.01, -0.001], *p* = .001), with no difference between the groups in Session 2 (^𝛽^^ = 0.001, 95% CI [-0.001, 0.004], *p* = .29). Analysis of trial time revealed a main effect of Block (*F*(1, 20.04) = 31.72, *p* <.001), however there were no significant interactions with Drug Condition or Session.

#### Retention

Assessment of offline change in skill revealed no main effect of Learning Session (*F(*1, 18) = 0.01, *p* = .91), nor was there a main effect of Drug Order (*F(*1, 18) = 1.69, *p* = .21). However, there was a significant interaction between Learning Session and Drug Order (*F(*1, 18) = 6.47, *p* = .02, 𝜂_p_^2^= .27). Bonferroni-corrected contrasts showed that participants who ingested sulpiride in session 1 showed greater offline improvement compared to placebo (*t*(35.54) = 2.66, *p* = .02, *d* = 1.42), with no difference in session 2 (*t*(35.54) = 2.66, *p* = .94).

The force error subcomponent of the skill measure showed no main effect of Learning Session (*F(*1, 18) = 0.48, *p* = .50), or Drug Order (*F(*1, 18) = 2.81, *p* = .11), however there was once again an interaction between Learning Session and Drug Order (*F*(1, 18) = 4.70, *p* = .04, 𝜂_p_^2^ = .21). Participants who ingested sulpiride in session 1 showed a reduction in force error at retention relative to placebo (*t*(31.31) = -2.60, *p* = .03, *d* = 1.15), with no difference between groups after session 2 (*t*(31.31) = -0.20, *p* >.99).

The trial time subcomponent of the skill measure showed a main effect of Learning Session (*F*(1, 18) = 5.56, *p* = .03, 𝜂_p_^2^ = .24), Drug Order (*F*(1, 18) = 6.06, *p* = .02, 𝜂_p_^2^ = .25), and a Learning 𝑝 𝑝 Session by Drug Order interaction (*F*(1, 18) = 4.40, *p* = .05, 𝜂_p_^2^= .20). Participants who ingested sulpiride in session 1 showed an increase in trial time, i.e. performance was slower, at the retention test, while those taking placebo did not (*t*(33.93) = 3.23, *p* = .01, *d* = 1.28). In contrast, there was no difference between sulpiride and placebo after session 2 (*t*(33.93 = 0.66, *p* >.99), as both groups performed slower at the retention test.

The duration of delay between acquisition and retention test did not differ between conditions *t*(39) = 1.34, *p =* .19). Furthermore, delay duration was not correlated with skill (ρ = - 0.003, 95% BCa CI [0.25, -0.31], *p* = .98), force error (ρ = -0.25, 95% BCa CI [0.41, -0.17], *p* = .39) or trial time (ρ = -0.28, 95% BCa CI [0.06, -0.53], *p* = .08) at retention.

## Discussion

The present study had three key findings. First, counter to expectation, 800 mg of sulpiride reduced performance during acquisition of a novel motor skill. However, performance improved significantly at the retention test once dopamine transmission was restored. Second, selective D2 blockade during acquisition resulted in slower performance at the retention test. Notably, these findings were specific to tasks learned in session 1. No difference was observed between sulpiride and placebo conditions on session 2 learning, i.e. when motoric demands differed, but the overall features of the task were familiar. Finally, we found no evidence that D2 blockade reduced offline consolidation. These results highlight the importance of D2 dopamine transmission in motor sequence learning and provide insight into the role of dopamine in online motor learning, particularly during initial learning when a task is unfamiliar.

### D2 blockade impaired motor performance during motor skill acquisition

Previous studies using sulpiride in healthy participants found no impact on processing speed, fine motor control (McClelland et al. 1990) or spatial working memory and planning (Mehta et al. 2003). Consistent with this literature, in the present study sulpiride did not impact the ability to perform high-intensity cycling exercise, pinch grip MVC, or baseline (block 1) motor performance. Sulpiride was also not expected to significantly impact online motor skill, but this is not what we observed. The present results highlight the importance of dopamine D2 transmission in the acquisition of a novel motor skill.

Conventional models of basal ganglia function propose that coordinated signalling along direct and indirect pathways allows for the selection and execution of desired motor output, and the simultaneous suppression of unwanted motor output (Freeze et al. 2013; Mink 2003). Supporting this model, studies in animals have shown that activation of both direct and indirect pathways is involved movement planning and execution (Cui et al. 2013; Nonomura et al. 2018). Specifically, activity of D2 indirect pathway neurons is associated with the initiation of a movement sequence (Augustin et al. 2020; Jin et al. 2014), supporting the role of D2 pathways in action selection and inhibition of undesired motor programs (Nonomura et al. 2018). Considered within this model, our finding that sulpiride reduced accuracy during the acquisition of a novel motor sequence task is likely the result from an imbalance between direct and indirect pathways in the associative and sensorimotor cortico-basal ganglia loops (Alexander and Crutcher 1990). The current study involved learning several distinct motor sequences (three per session) which had to be selected and executed rapidly according to the target locations. The selective blockade of D2 receptors may have disrupted the suppression of the alternative motor sequences (Calabresi et al. 2014; Mink 1996; Redgrave et al. 2010), thereby impacting selection of the appropriate sequence of force pulses to reach the targets in a given trial.

In addition to an effect of D2 blockade on sequence selection and initiation, sulpiride may have impacted precision force generation. Functional magnetic resonance imaging (fMRI) studies implicate the basal ganglia in precision grip force control (Prodoehl et al. 2009). Specifically, activity in anterior basal ganglia nuclei is associated with planning grip force (Wasson et al. 2010), while posterior basal ganglia nuclei support on-line adjustments during precision grip (Grafton and Tunik 2011). This is supported by studies in PD, as disruption in precision grip is a common motor symptom and has been linked to disease severity (Fellows et al. 1998; Roberts et al. 2015). PET imaging studies have demonstrated reduced dopamine uptake and receptor density in the striatum of PD patients compared to controls (Brück et al. 2006; Ouchi et al. 2005). These metabolic changes are consistent with an fMRI study by Spraker et al. (2010) showing reduced activity in the basal ganglia of PD patients during a force grip task. Notably, reduced activation was more prominent with increased rapid switching between contraction and relaxation. Disruption of action selection mechanisms have been proposed as a key mechanism underlying motor symptoms in disorders of dopamine depletion such as PD (Wiecki and Frank 2010). The current findings provide support for the importance of D2 dopamine in action selection during motor skill learning, specifically in tasks requiring precision grip.

Importantly, differences in acquisition were only observed during session 1, with no difference between sulpiride and placebo in session 2 when a different force-cursor transformation and set of sequences were acquired. This may be explained by overall familiarity with general task requirements. Session 1 learning likely involved additional explicit learning requirements in addition to learning the specific motor sequences, such as memorising when to start, the target order (red-blue-green-yellow-white), etc. Parallel learning processes, i.e. explicit and implicit learning, can interfere with one another (Albert et al. 2022), and the addition of explicit learning demands may have exacerbated the impact of dopamine D2 blockade on motor performance, and increased task difficulty, in the first session. This is supported by previous studies investigating the impact of sulpiride on spatial planning and working memory, which found that deficits associated with sulpiride increased with task difficulty (Naef et al. 2017). Participants who ingested sulpiride in session 2 had prior exposure to the task while on placebo, which may have conferred some protection from the impact of sulpiride (Mehta et al. 1999). This suggests that D2 blockade has a specific impact on the performance of novel motor tasks with greater explicit learning demands.

Task performance may have been impacted by general sedative effects of sulpiride on sensorimotor processing rather than drug specific mechanisms on motor learning. However, subjective alertness ratings did not differ across drug conditions or sessions. Indeed, participants in the current study reported increased alertness following exercise, immediately prior to completing the motor task. The potential sedative effects of sulpiride were evidently counteracted by the invigorating effects of exercise, at least at the beginning of the task (Dietrich and Audiffren 2011). This suggests sulpiride-induced reductions in performance during motor skill learning cannot be explained by a sedative effect of sulpiride.

### The impact of exercise and D2 blockade on retention of motor skill

In addition to facilitating motor output, dopamine is instrumental in supporting motor learning by modulating cortical plasticity (Macedo-Lima and Remage-Healey 2021), which underlies motor consolidation. Further, acute exercise has been shown to improve dopamine-mediated plasticity and motor learning and may act via D2-related mechanisms (Christiansen et al. 2019; Mang et al. 2017). It was anticipated that blockade of D2 dopamine would disrupt consolidation of a motor skill, resulting in poorer performance at a 7-day retention test compared to placebo. It was further expected that acute exercise would amplify this effect, by enhancing learning in the placebo group. Contrary to expectation, there was no impact of sulpiride on motor consolidation. There was a difference in offline learning between sulpiride and placebo conditions following session 1, with the sulpiride group showing an offline increase in skill between initial learning and retention, resulting in similar performance to placebo at the retention test.

Given the impact of sulpiride on the acquisition phase of learning, it is likely any impacts of dopamine D2 blockade and high intensity exercise on consolidation were masked by the absence of drug at the retention test. However, these results are still informative. Similar skill performance for both groups at retention, when both groups completed the task without drug, indicates learning took place during acquisition in both placebo and sulpiride conditions. Notably, the groups differed in the speed and accuracy subcomponents of skill during session 1 retention, with placebo participants improving speed at the cost of accuracy, while sulpiride participants showed greater accuracy but performance was slowed. Motor learning is supported by dopamine-driven reward and error signals within the basal ganglia (Festini et al. 2016; Galea et al. 2015; Schultz et al. 1997), with reward increasing speed and movement vigour (Codol et al. 2020; Galea et al. 2015). As successful completion of a motor skill is inherently rewarding (Badami et al. 2012; Blain and Sharot 2021), greater accuracy in the placebo group during acquisition may have led to increased speed at retention. In contrast, participants who learned on sulpiride utilised a slower approach to compensate for reduced accuracy during acquisition (Fitts 1954; Thumser et al. 2018). This is consistent with studies indicating D2 antagonism may increase sensitivity to negative outcomes (Hong and Hikosaka 2011; Palminteri et al. 2011), and result in greater reliance on avoidance-based learning (Baetu et al. 2015; Wiecki and Frank 2010).

### Limitations

The impact of sulpiride was specific to the motor sequences learned in session 1, which limited comparisons between sessions 1 and 2 and reduced the overall power for analysis of motor learning retention. However, it is unlikely that the similar performance across groups in session 2 reflects ceiling effects, as previous studies have demonstrated improvement in SVIPT tasks across training sessions and over multiple days (Cantarero et al. 2013; Reis et al. 2009). Furthermore, comparison of performance during retention tests was confounded by the effect of sulpiride during acquisition, as the retention test was conducted without the drug. However, a benefit of the present study’s within-subjects design was the ability to assess the differential impact of D2 blockade on novel motor sequence learning versus when some general task features were familiar.

### Conclusions

We show that sulpiride impaired accuracy during the acquisition of a novel motor skill following exercise, however participants were able to demonstrate learning once normal dopamine processing was restored. Additionally, sulpiride impacted how the task was approached upon retention, resulting in a slower performance to prioritise accuracy. This highlights the role of the dopamine D2 receptor on motor tasks requiring precision force control and may have functional relevance in motor rehabilitation in disorders such as PD, by informing the timing of motor learning in conjunction with dopaminergic medication. For tasks requiring fine motor control and precision grip, aberrant dopamine functioning may have a more prominent impact when tasks are unfamiliar or difficult. Timing novel learning with peak efficacy of dopaminergic medication may improve learning outcomes in the longer term.

## Acknowledgements

We thank Dr Ziarih Hawi, Daniel Pearce, and Julia Koutoulogenis for their assistance with drug holding and blinding; Claire Cadwallader and Sarah Cohen for their support with data collection; and Bridgitt Shea, Amy Huynh, Huw Jarvis, and Dami Obawede for their assistance with participant recruitment.

## Authorship contribution statement

**Eleanor M. Taylor:** Conceptualization, Methodology, Formal analysis, Investigation, Writing – Original Draft, Visualisation. **Dylan Curtin:** Conceptualization, Methodology, Formal analysis, Investigation, Writing - Review & Editing. **Mark A. Bellgrove:** Conceptualization, Methodology, Writing - Review & Editing, Funding acquisition. **Trevor T-J. Chong:** Conceptualization, Methodology, Writing - Review & Editing, Funding acquisition. **James P. Coxon:** Conceptualization, Methodology, Formal analysis, Investigation, Writing - Review & Editing, Visualisation, Funding acquisition.

