## Supplementary Materials for "The effect of dopamine D2 receptor blockade on human motor skill learning"

**Supplementary Material: "Dopamine D2 receptor blockade reduces motor skill learning in healthy young adults"**

**Contents**

**Supplementary Methods (pp. 2-5)**

Exercise protocol

Linear mixed model fitting

Supplementary Table 1: Model fitting for skill measure

Supplementary Table 2: Model fitting for force error

Supplementary Table 3: Model fitting for trial time

**Supplementary Results (pp. 5-6)**

Drug and exercise characteristics

Supplementary Table 4: Summary of exercise performance

Supplementary Table 5: Summary of LMM results

**Supplementary Figures (pp. 7-9)**

Supplementary Figure 1: Participant recruitment flow diagram

Supplementary Figure 2: Bond-Lader visual analogue scale (VAS) ratings as a function of drug condition

### Supplementary Methods

#### Exercise Protocol

The exercise protocol was completed on a stationary Wattbike Atom exercise bike (Wattbike, 2017). To ensure equivalent relative intensity across participants, exercise was tailored to each individual based on their estimated heart rate reserve (HRR):

$$HRR = (HR_{age-predicted\ max} - RHR)$$

where

$$HR_{age-predicted\ max} = 208 - (0.7 * age)$$

{Tanaka, 2001 #495}

The 20-minute cycling protocol alternated between 3-minute phases of low- to moderate-intensity cycling (approximately 50% HRR) and 2-minute phases of high-intensity cycling (up to 90% HRR). This was followed by a brief low-intensity cool-down period.

#### Linear Mixed Model Fitting

The Linear Mixed Models (LMM) for the current study were constructed using a model selection approach. A maximal model-based approach was considered inappropriate for the current study, given the small sample size and risk of overfitting {Matuschek, 2017 #1087}. Variables of theoretical interest were entered into the model and were retained if they significantly improved overall model fit as indicated by the Akaike Information Criterion (AIC) {Meteyard, 2020 #1089}. In cases where chi-square comparisons between models were non-significant, the more parsimonious model was selected. See Table S1 for a summary of model comparisons. Additional models were constructed using the above criteria to assess force error and trial time (See Table S2 and S3). Models were fit in RStudio (version 2022.07.2) (RStudio Team, 2022) using the lme4 package {Bates, 2015 #1090}. Model comparisons were conducted with maximum likelihood estimation, with restricted maximum likelihood (REML) estimation and a Satterthwaite adjustment to compute the degrees of freedom in the final models. See Table S4 for a summary of selected models. Overall effects, interactions and *p* values were calculated using the lmerTest package {Kuznetsova, 2017 #1093}. Effect sizes were computed as outlined by Brysbaert and Stevens {Brysbaert, 2018 #1097}.

**Table S1.** *Model fitting for skill measure.*

| Sampling Units | | N total obs = 528; N Subjects = 22; N blocks = 12; N Sessions = 2 | | | | | | | | |
|---|---|---|---|---|---|---|---|---|---|---|
| **Model Specification** | **Model Name** | **Compa-rison Model** | **Model description** | | **Random Effects^** | **Model Fit** | | | **LRT Test against comparison** | |
| | | | | | | **AIC** | **BIC** | **LL** | $\chi^2$ | *p* |
| RE only | Null | | Skill ~ (+1 \| Participant) | | - | -353.28 | -340.49 | 179.64 | | |
| | Skill 1 | Null | Skill ~ (1\|Block) +(1\|Participant) | | Block | -403.99 | -386.94 | 206.00 | 52.71 | <.001* |
| | Skill 2 | Skill 1 | Skill ~ (Block \| Participant) | | Block | -406.90 | -385.58 | 208.45 | 4.91 | .03* |
| FE main effects | Skill 3 | | Skill ~ Block + (Block \| Participant) | | Block | -432.51 | -406.93 | 222.26 | 27.61 | <.001* |
| | Skill 4 | Skill 3 | Skill ~ Block + Drug + (Block \| Participant) | | Block | -455.21 | -425.37 | 234.61 | 24.70 | <.001* |
| | Skill 5 | Skill 4 | Skill ~ Block + Drug + Session + (Block \| Participant) | | Block | -550.19 | -516.09 | 283.10 | 96.98 | <.001* |
| FE Interactions | Skill 6 | Skill 5 | Skill ~ Block * Drug + Session + (Block \| Participant) | | Block | -553.70 | -515.33 | 285.85 | | |
| | **Skill 7** | **Skill 6** | **Skill ~ Block * Drug * Session + + (Block \| Participant)** | | **Block** | **-560.64** | **-509.48** | **292.32** | **115.43** | **<.001*** |

Final selected model in bold.
^participant was included as a random grouping effect in all models
*$p$ <0.5

**Table S2.** *Model fitting for force error.*

| Sampling Units | | N total obs = 528; N Subjects = 22; N blocks = 12; N Sessions = 2 | | | | | | | | |
|---|---|---|---|---|---|---|---|---|---|---|
| **Model Specification** | **Model Name** | **Compa-rison Model** | **Model description** | | **Random Effects included^** | **Model Fit** | | | **LRT Test against comparison** | |
| | | | | | | **AIC** | **BIC** | **LL** | $\chi^2$ | *p* |
| RE only | Null | | Error ~ (+1 \| Participant) | | | -1460.3 | -1447.3 | 733.05 | | |
| | Error 1 | Null | Error ~ (1\|Block) +(1\|Participant) | | Block | -1467.6 | -1450.5 | 737.80 | 9.50 | .002* |
| | Error 2 | Error 1 | Error ~ (Block \| Participant) | | Block | -1488.6 | -1467.2 | 749.28 | 22.97 | <.001* |
| FE main effects | Error 3 | Error 2 | Error ~ Block + (Block \| Participant) | | Block | -1495.4 | -1469.8 | 753.70 | 8.84 | .003* |
| FE main effects | Error 4 | Error 3 | Error ~ Block + Drug + (Block \| Participant) | | Block | -1495.6 | 1465.7 | 754.79 | 2.17 | .14 |
| | Error 5 | Error 4 | Error ~ Block + Drug + Session + (Drug \| Participant) | | **Block** | -1524.6 | -1490.5 | 770.29 | 31.00 | <.001 |
| FE Interactions | Error 6 | Error 5 | Error ~ Block * Drug + Session + (Drug \| Participant) | | **Block** | -1527.6 | -1489.2 | 772.81 | 5.05 | .03* |
| | **Error 7** | **Error 8** | **Error ~ Block * Drug * Session * (Drug \| Participant)** | | Block | **-1533.5** | **-1482.3** | 778.76 | **11.89** | **.01 *** |

Final selected model in bold.
^participant was included as a random grouping effect in all models
**p* <0.5

**Table S3.** *Model fitting for trial time.*

| Sampling Units | N total obs = 528; N Subjects = 22; N blocks = 12; N Sessions = 2 | | | | | | | | | |
|---|---|---|---|---|---|---|---|---|---|---|
| **Model Specification** | **Model Name** | **Compa-rison Model** | **Model description** | | **Random Effects included^** | **Model Fit** | | | **LRT Test against comparison** | |
| | | | | | | **AIC** | **BIC** | **LL** | $\chi^2$ | *p* |
| RE only | Null | | Time ~ (1 \| Participant) | | | 711.51 | 724.30 | -352.76 | | |
| | Time 1 | Null | Time ~ (1\|Block) + (1\|Participant) | | Block | 680.54 | 697.59 | -336.27 | 32.98 | <.001* |
| | Time 2 | Time 1 | Time ~ (Block \| Participant) | | Block | 667.60 | 688.91 | -328.80 | 14.94 | <.001* |
| FE main effects | Time 3 | Time 2 | Time ~ Block + (Block \| Participant) | | Block | 650.41 | 675.99 | -319.21 | 19.18 | <.001* |
| | Time 4 | Time 3 | Time ~ Block + Drug + (Block \| Participant) | | Block | 582.35 | 612.20 | -284.18 | 70.06 | <.001* |
| | Tine 5 | Time 4 | Time ~ Block + Drug + Session (Block \| Participant) | | Block | 381.31 | 415.42 | -182.65 | 203.04 | <.001* |
| FE Interactions | **Time 6** | | **Time ~ Block * Drug * Session + (Drug \| Participant)** | | **Block** | **384.64** | **435.80** | **-180.32** | | |

Final selected model in bold.
^participant was included as a random grouping effect in all models
*$p$ <0.5

### Supplementary Results

#### Drug and Exercise Characteristics

***Side Effects.*** Four participants reported fatigue and restlessness several hours after the Sulpiride testing session, and two participants reported fatigue following Placebo. All side effects were fully reversible, did not necessitate treatment, and did not interfere with participants' ability to complete the required tasks. No further adverse events were reported.

A LMM was used to compare subjective ratings of alertness, calmness, and contentedness in the Sulpiride and Placebo conditions across the four VAS time points. Session (1 and 2), Drug Condition (Sulpiride and Placebo), and Timepoint (T0, T1, T2, T3) were entered as fixed effects, with Participant entered as a random effect. No significant main effects or interactions were seen for alertness or contentedness ratings (all $p$ >.05). Subjective calmness ratings were lower at T3 (immediately following exercise) compared to T0 ($t(136) = 5.18$, $p$ <.001, $d$ = .71), however this did not differ significantly across Drug condition ($t(136) = -0.010$, $p$ = .92) or Session ($t(136) = -1.66$, $p$ =

.09), and likely reflects the anxiolytic effects of high intensity exercise. A visual depiction of VAS ratings by Drug Condition and Timepoint is shown in Figure S1.

***Evaluation of Participant Blinding***. A subset of participants (N = 13) were polled about which session they thought they took Sulpiride vs. Placebo. Fisher's exact test, which is recommended for small samples ($N < 20$; Kim, 2017) revealed no significant differences in the proportion of people who correctly vs incorrectly guessed the drug sequence order ($P = 0.56$), indicating that participant blinding was maintained.

**Table S5**

*Summary of Exercise Performance Across Placebo and Sulpiride Conditions*

| Exercise measure | Placebo | Sulpiride | Test | *p* |
|---|---|---|---|---|
| Average cadence (RPM) | 75.74 (6.22) | 75.40 (6.22) | $t(21) = 0.84$ | 0.41 |
| Average power (Watts) | 91.85 (22.29) | 90.78 (21.30) | $t(21) = 0.84$ | 0.41 |
| Average power:weight (Watts/Kg) | 1.25 (0.22) | 1.27 (0.24) | $t(21) = 0.96$ | 0.35 |
| Peak heart rate reserve, % | 90% (7.91) | 92% (8.39) | $t(18) = 1.00$ | 0.33 |
| Peak cadence (RPM) | 112.45 (16.79) | 110.14 (14.82) | $t(21) = 1.02$ | 0.32 |
| Peak power (Watts) | 285.55 (111.73) | 284.86 (105.45) | $t(21) = 0.05$ | 0.96 |
| Peak power:weight (Watts/Kg) | 3.89 (1.10) | 3.90 (1.17) | $t(21) = 0.08$ | 0.93 |
| Peak Perceived Exertion (Borg) | 17.23 (2.25) | 17.41 (1.89) | $t(21) = 0.78$ | 0.45 |

Values are means (standard deviation). RPM = revolutions per minute.

***Exercise Analysis***. Heart rate, cadence, and power (Watts) were collected and analysed across 1-second intervals. RPEs were collected each minute. Heart rate data from 3 sessions was not collected due to technical difficulties with the heart rate monitor. A summary of exercise details is outlined in Table S5.

**Table 3**
Results of linear mixed models for acquisition across skill, force error and trial time measures

| Parameter | Estimate (*SE*) | 95% CI [lower, upper] | Test *(df)* | *p* |
|---|---|---|---|---|
| | | **Skill** | | |
| Drug (reference = placebo) | 0.34 (0.05) | 0.24, 0.43 | *F*(1, 477.59) = 0.31 | .58 |
| Block | 0.03 (0.003) | 0.02, 0.04 | *F*(1, 20.10) = 74.84 | **<.001*** |
| Session (reference = Session 1) | 0.13(0.01) | -0.01, 0.27 | *F*(1, 477.59) = 45.12 | **<.001*** |
| Drug × Block | -0.02 (0.01) | -0.03, -0.01 | *F*(1, 477.52) = 5.53 | **.02*** |
| Placebo vs Sulpiride | 0.003 (0.002) | 0.00, 0.01 | *t*(478) = 2.35 | **.02*** |
| Drug × Session | 0.07 (0.13) | -0.19, 0.32 | *F*(1, 20.04) = 0.25 | .62 |
| Block × Session | -0.02 (0.01) | -0.03, -0.01 | *F*(1, 477.52) = 4.33 | **.04*** |
| Session 1 vs Session 2 | 0.003 (0.002) | 0.00, 0.01 | *t*(478) = 2.08 | **.04*** |
| Drug × Block × Session | 0.02 (0.01) | 0.01, 0.04 | *F*(1, 20.10) = 8.83 | **.01*** |
| Drug × Block trend | | | | |
| Session 1 | 0.01 (0.003) | 0.004, 0.02 | *t*(53.9) = 3.79 | **<.001*** |
| Session 2 | -0.002 (0.003) | -0.01, 0.004 | *t*(53.8) = -0.83 | .41 |
| | | **Force Error** | | |
| Drug (reference = placebo) | -0.01 (0.03) | -0.06 – 0.05 | *F*(1, 477.32) = 1.60 | .21 |
| Block | -0.01 (0.002) | 0.00, 0.01 | *F*(1, 19.95) = 14.14 | **.001*** |
| Session (reference = Session 1) | -0.03 (0.03) | -0.09 – 0.02 | *F*(1, 477.32) = 18.94 | **<.001*** |
| Drug × Block | 0.01 (0.002) | 0.004 – 0.01 | *F*(1, 477.26) = 5.05 | **.03*** |
| Placebo vs Sulpiride | | | | |
| Drug × Session | 0.01 (0.05) | -0.11, 0.08 | *F*(1, 20.02) = 0.07 | .79 |
| Block × Session | 0.01 (0.002) | 0.003 – 0.01 | *F*(1, 477.26) = 3.68 | .06 |
| Drug × Block × Session | -0.01 (0.004) | -0.02, -0.003 | *F*(1, 19.95) = 8.13 | **.01*** |
| Drug × Block trend | | | | |
| Session 1 | -0.004 (0.001) | -0.01, -0.001 | *t*(42.8) = -3.62 | **.001*** |
| Session 2 | 0.001 (0.001) | -0.001, 0.004 | *t*(42.7) = -1.08 | .29 |
| | | **Trial Time** | | |
| Drug (reference = placebo) | 0.52 (0.37) | -0.19 – 1.24 | *F*(1, 477.21) = 29.68 | **<.001*** |
| Block | -0.03 (0.01) | -0.06, -0.01 | *F*(1, 20.04) = 31.72 | **<.001*** |
| Session (reference = Session 1) | -0.26 (0.36) | -0.98 – 0.45 | *F*(1, 477.21) = 74.56 | **<.001*** |
| Drug × Block | -0.03 (0.02) | -0.06, 0.01 | *F*(1, 477.19) = 0.21 | .65 |
| Drug × Session | -0.44 (0.72) | -1.86 – 0.98 | *F*(1, 19.99) = 0.37 | .55 |
| Block × Session | -0.01 (0.02) | -0.05, 0.02 | *F*(1, 477.19) = 1.84 | .18 |
| Drug × Block × Session | 0.05 (0.03) | -0.01 – 0.10 | *F*(1, 20.04) = 2.45 | 0.13 |

Df = degrees of freedom. *F* statistic derived from type III ANOVA. Pairwise comparisons and trends are indented. **p* >.05

**Supplementary Figures**

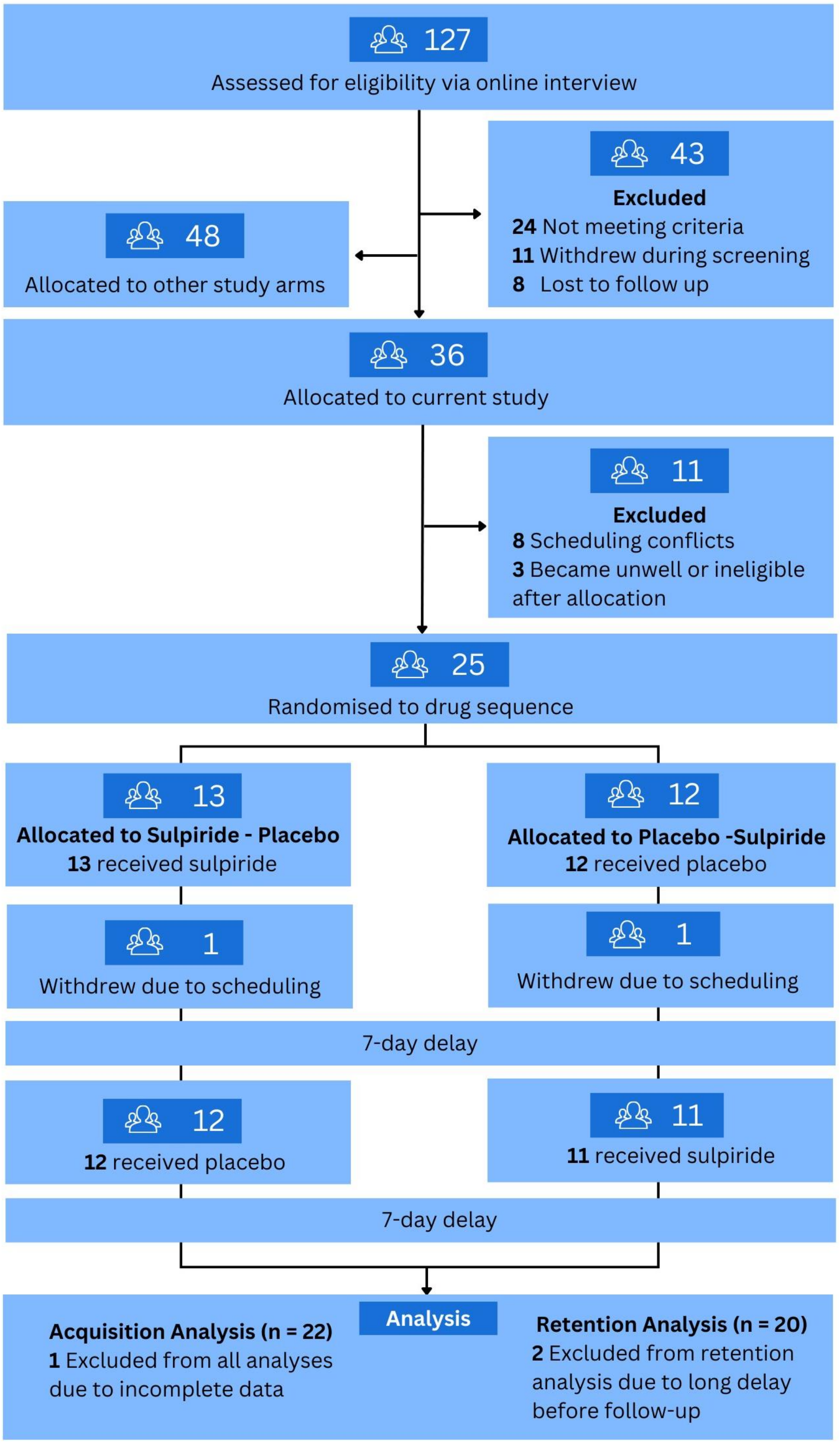


**Figure S1. Participant inclusion and attrition across our cross randomised, double-blind, crossover trial**. Participants were recruited via online notices and community flyers, with data was collected from June 2021 to August 2022 at Monash University, Melbourne.

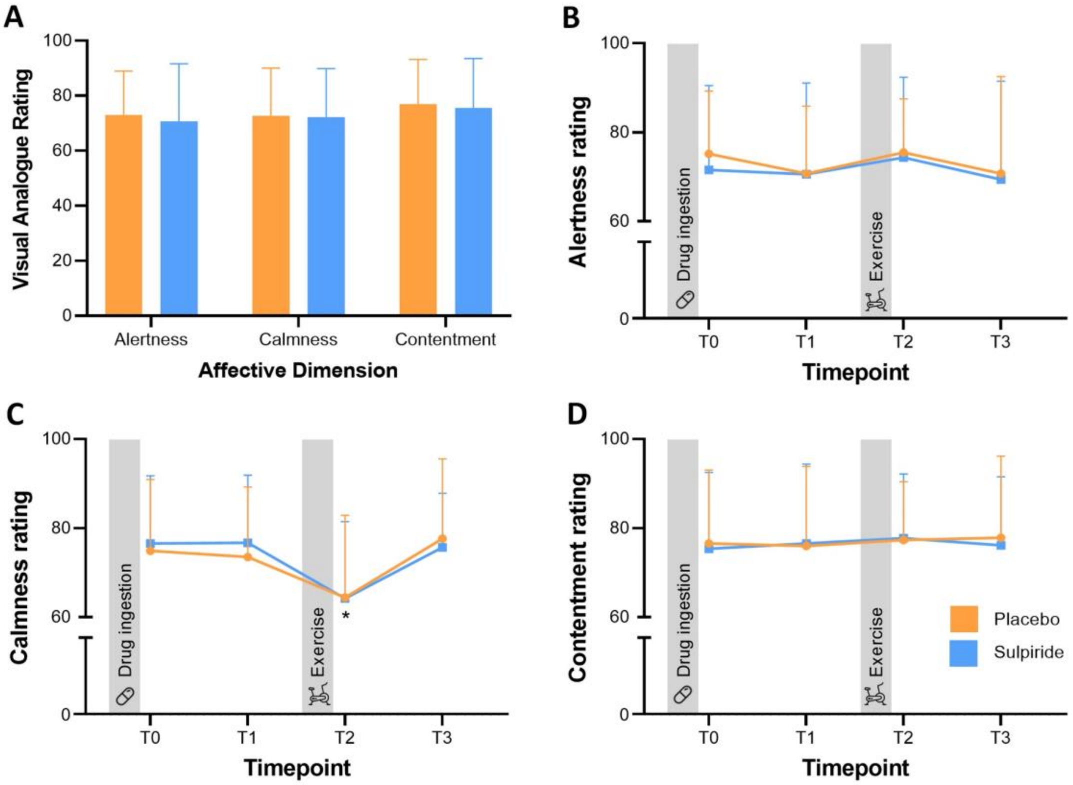


**Figure S2**. Bond-Lader Visual Analogue Scale (VAS) Ratings Across Placebo and Sulpiride Sessions. Means and standard deviations are shown for the three affective dimensions of the scale (alertness, calmness, contentment). Higher scores (range 0-100) denote higher positive affect. (A) Average ratings for each affective dimension across all timepoints. (B-D) Mean ratings for each dimension across timepoints. The first rating (T0) was taken immediately after drug ingestion, with subsequent ratings taken every hour thereafter. Calmness was rated lower at T2, immediately after exercise, however there was no difference between drug conditions. Alertness and contentment did not differ significantly across timepoints.

*$p$ <.05
